# The effect of climate change on the spread of predicted bluetongue in Australian livestock

**DOI:** 10.1101/2024.05.01.592030

**Authors:** S Al-Riyami, SM Firestone, D Eagles, R Bradhurst, MA Stevenson

## Abstract

**Introduction:** The frequency of vector-borne disease in human and animal populations has increased in recent years leading to concerns that even greater increases will occur as a result of climate change, driven by changes in the geographic distribution of insect vector habitat areas. In this study we investigate the effect of climate change on the expected spread of bluetongue (a viral disease of ruminants spread by *Culicoides* midges) using the Australian Animal Disease Spread model (AADIS).

**Methods:** Estimates of average daily temperature across Australia for 2015 were obtained from the Australian Bureau of Meteorology. Predicted average daily temperatures using the CanESM2 model (emission scenario RCP 8.5) for 2025 and 2035 at a resolution of 5 km × 5 km were obtained from the Australian Climate Futures decision-support tool, managed by CSIRO. Two study areas in Australia were selected: the first in North Queensland and the second in Northern New South Wales. A total of 24 outbreak scenarios were run: mid-summer and mid-winter incursions for each study area for 2015, 2025 and 2035 with direct movement of animals in the AADIS model disabled and enabled for each. Model results were expressed as the number bluetongue-positive herds, (herds in which at least one animal was BTV positive) at the end of each 365 day simulation period.

**Results:** For North Queensland, there was little change in the median predicted number of bluetongue positive herds for mid-summer and mid-winter 2025 and 2035 incursions (compared with 2015) and a moderate increase in the variability of predicted outbreak sizes when direct animal movements were disabled. For Northern New South Wales there were moderate increases in both the predicted number of bluetongue positive herds and the variability of predicted outbreak sizes for 2025 and 2035, compared with 2015. Compared with the direct animal movement disabled scenarios, there were marked increases in the predicted number of bluetongue positive herds as a function of simulation year for North Queensland. For Northern New south Wales this trend was not as distinct, but as for the direct movement disabled scenarios, the variability of predicted outbreak sizes for the 2035 incursions were greater than the variability of predicted outbreak sizes for the 2015 incursions.

**Conclusion:** Climate change will result in a greater portion of the land area of Australia with conditions suitable for *Culicoides* midges. Our findings show that under conditions of climate change and an outbreak of virulent bluetongue in Australia, the rapid imposition of effective restrictions of animal movement will be the single most important control measure to limit further spread of disease.

## Introduction

Climate change is considered to be one of the greatest threats to human health by the World Health Organization (1). The environmental consequences of climate change which include sea-level rise, increased frequencies of flood and drought, more intense hurricanes and storms, heat waves and degraded air quality are predicted to have substantial impacts on human well-being (2, 3).

Over the past few decades, global warming has caused pronounced and often complex changes in the incidence and prevalence of vector borne infectious diseases of humans and animals (4, 5, 6). Following the emergence and re-emergence of vector-borne diseases (VBD) of humans in many parts of the world in recent years (e.g., Zika virus and malaria, Japanese Encephalitis) the global spread of arboviruses has received research attention from international public health organisations (7, 8, 9). There is a strong association between climate change and the geographic distribution of VBD because insect vector ecology and their rate of development are often dependent on environmental conditions. Generally, larval development stages are dependent on the presence of bodies of water and/or specific conditions of humidity and temperature. Vector biting rates are positively associated with temperature up to a species-specific threshold (10, 11). The amount of time taken for pathogens to replicate within insect vectors (the extrinsic incubation period) is known to decrease when temperatures are higher (12, 13) and virus survival times are increased with increases in environmental temperatures (10).

Given the sensitivity of arthropods to changes in climatic conditions it is reasonable to expect that climate change will result in marked changes in the geographic extent of VBD and marked changes in the seasonality of their occurrence (14). In human medicine, studies estimating the changes in the geographic distribution of malaria has recently received the bulk of research attention at both the global (15, 16) and regional level (17, 18). Yet, a lesser number of studies have addressed the importance of socioeconomic factors (16, 19, 20), economic development and interventions which might be developed to offset or counteract climate-induced trends (15). These studies are unable to account for the influence of other drivers of disease for two reasons: (1) there is often little or no quantitative evidence to define how the frequency of disease will be affected by change in a non-climate driver; and (2) there are few models developed to predict how these non-climatic drivers could change in the future. A 2019 study assessed changes in vector abundance, factors influencing vector population dynamics, host density and land use, and concluded that the changes in disease frequency that were observed were likely due to observed changes in climate (21). This conflicts with the relative simplicity of climate disease modelling where observational or experimental data are used to associate weather or climate variables with the frequency of disease. Combining the two allows climate sensitive disease patterns or distributions to be predicted under future climate scenarios. Bluetongue (BT), a viral disease of ruminants causing fever, lameness and swelling of the lips and tongue, is likely to be one of the most important diseases of animals influenced by climate change due to expected changes in the geographic distribution of *Culicoides* midges (22). In Europe, a series of BT outbreaks occurred after 1998 (23) resulting in the death of millions of ruminants and substantial economic losses. Between 1998 and 2010 more than 110,000 BT outbreaks in Europe were reported to the World Organisation for Animal Health. Over 80,000 of these were due to bluetongue virus (BTV) serotype 8. In 2006, bluetongue occurred for the first time in northern Europe (24). In 2007 bluetongue spread to the United Kingdom most likely via wind-borne infected midges blown across the English Channel from mainland Europe (25). In 2015, bluetongue re-emerged in France (26). An outbreak of BTV-4 that started in the French Alps (in the southeast of the country) in 2017 had spread as far as Normandy (in the far north west) by 2018 (27).

Expansion of the geographic distribution of bluetongue in southern Europe has been hypothesised to be associated with change in the geographic extent of the Afrotropical midge vector (28). It is possible that climate has played a role in both routes of expansion (7, 29, 30). Several disease simulation models have been developed for bluetongue, all of which account for the effect of environmental temperature on the rate and geographic distribution of disease spread (31, 32, 33, 34). While these models provide an opportunity to quantify the effect of climate change on the epidemiology of bluetongue, predictions are likely to be imprecise due to uncertainties in climate change projection data and uncertainties in factors driving the epidemiology of the disease itself (35, 36, 37). This said, the results of modelling, if based on a valid conceptual design, provide useful information to guide animal health decision making (38). Bluetongue virus is already present in parts of Australia and, at the time of writing, the strains that are present are less virulent than strains that occur overseas (39). There is a risk that a novel or virulent BTV strain may be introduced into the country which would then have a substantial impact on animal health. To investigate this risk in detail the objectives of this study were to: (1) use climate change projection data to simulate the spread of bluetongue in Australia using the Australian Animal Disease Spread model (AADIS) (40, 41); and (2) provide recommendations on the way surveillance and outbreak management plans for bluetongue should be changed in the short to medium term in response to climate change.

## Materials and methods

### Bluetongue transmission model

The methodology involves simplifications and assumptions to model vector population biology using the AADIS model. Geographic automata were used to simplify population distribution into a gridded format, ignoring fine-scale variations within cells and between neighboring cells with differing states. Vector populations within each grid cell were represented on a relative scale from 0 to 1, based on a limiting resource, typically cattle density for certain species like the *culicoides* vector (42). The model does not explicitly represent all lifecycle stages due to computational constraints; instead, only adult vectors are modeled directly, with immature stages considered implicitly under unsuitable conditions. We configured AADIS with the *Culicoides* spp. densities distribution, abundance and bluetongue transmission model as described Al Riyami (2020). Population dynamics within each cell are modeled using a logistic growth equation, adjusted by the cell’s cattle density and temperature variations affecting vector recruitment and mortality rates (43). The model is designed to be flexible, allowing for changes in the growth equation based on new research or specific vector data. This adaptability is crucial for maintaining the model’s relevance and accuracy in different ecological contexts or in response to new findings.

Predictions of the spatial and temporal spread of bluetongue following introduction of the virus into single herd locations in 2015 provided a reference from which bluetongue spread scenarios under conditions of climate change (2025 and 2035) could be compared. BTV was seeded into two geographically distinct locations in Australia during the mid-summer (1 January) and mid-winter (1 July): (1) Mareeba at the base of Cape York Peninsula in North Queensland (labelled ‘A’ in Figure 1); and (2) Heron’s Creek which borders Kempsey in Northern New South Wales (labelled ‘B’ in Figure 1). We selected North Queensland as a study area because it is well known to be within the BTV transmission zone defined by the National Arbovirus Monitoring Program (NAMP) (National Arbovirus Monitoring Program 2015). Northern New South Wales was selected because of its location within the BTV transmission zone and the conditions favorable to the insect vector are observed seasonally, extending from north to south along the east coast of New South Wales (NSW) in most years (44).

**Figure 1:**
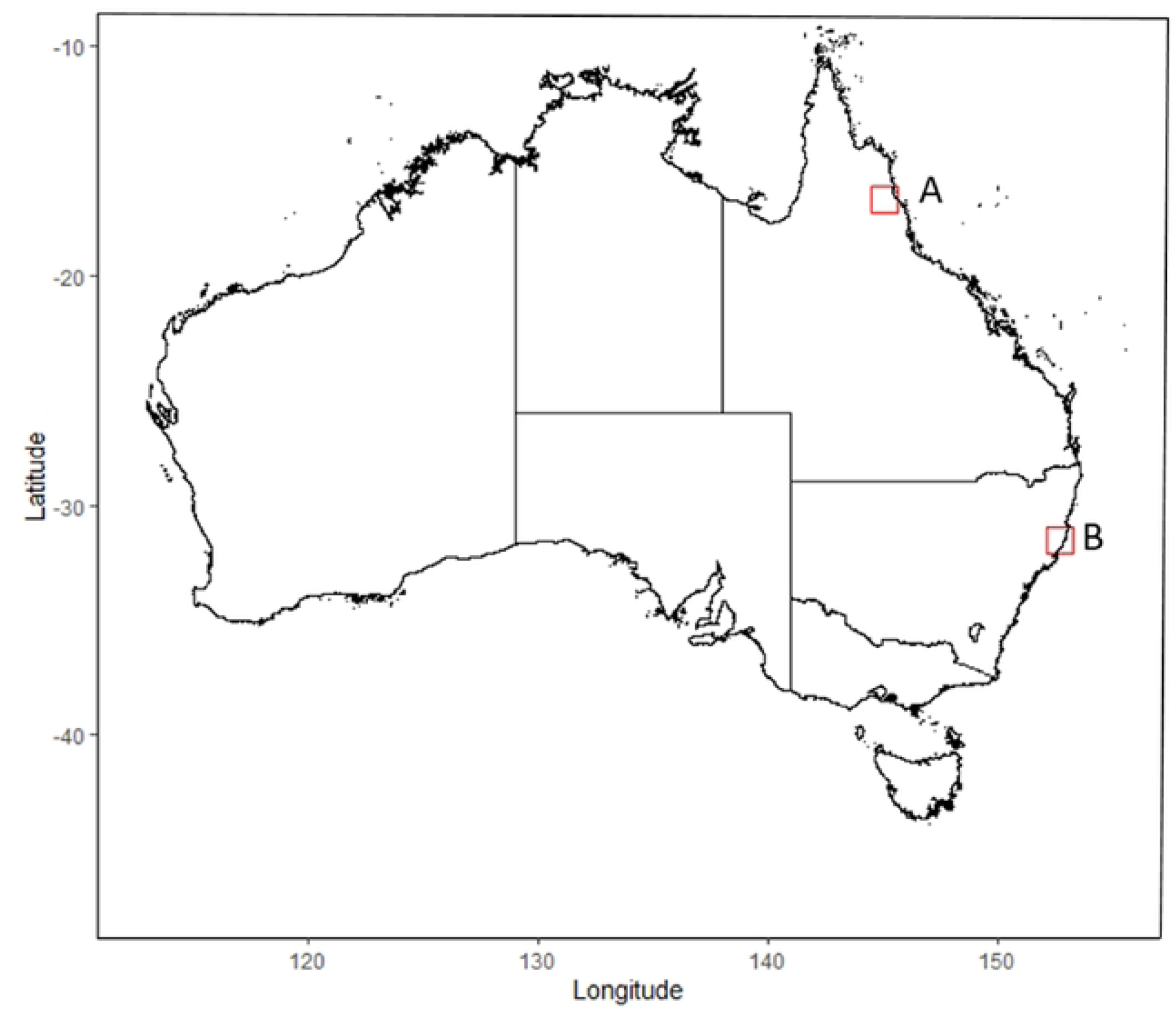
The effect of climate change on the spread of bluetongue in Australian livestock. Map of Australia showing the location of the study areas: (A) North Queensland, (B) Northern New South Wales.

### Climate model data

Climate change involves long-term alterations in weather patterns and changes in the physical characteristics of oceans, land, and ice sheets, driven by natural and anthropogenic factors, including the emissions of greenhouse gases like CO2 and CH4 (45). Since 2009, atmospheric carbon emissions have risen by approximately 2.0 ppm annually, predominantly due to industrial activities (46, 47). The phenomenon is linked to an energy imbalance caused by solar radiation absorption and re-emission by the Earth, with greenhouse gases trapping heat and influencing the hydrologic cycle, leading to varied global climate effects. The development of climate change scenarios has evolved from focusing on greenhouse gas emission factors to integrating emissions with socioeconomic pathways, known as Representative Concentration Pathways (RCPs) (48). These RCPs, ranging from very low (RCP 2.6) to very high (RCP 8.5) emission scenarios, provide a spectrum of future climate projections, factoring in greenhouse gas mitigation potentials without presuming the likelihood of any single scenario (48).

The future climate cannot be predicted on the basis of historical climate observations since it depends on future greenhouse gas emissions. Global climate models (GCMs) are generally regarded as the best tools for climate change projections (CSIRO and Bureau of Meteorology 2015). GCMs represent the atmosphere and bodies of water as a three-dimensional grid, with a typical atmospheric resolution of around 200 km and 20 to 50 levels in the vertical direction (CSIRO and Bureau of Meteorology 2015). These models are able to represent large-scale synoptic features of the atmosphere such as the development of high- and low-pressure systems and large-scale oceanic currents. Important physical processes occur at finer spatial scales including precipitation, cloud development and atmospheric and oceanic turbulence. The factors that influence these processes are included in ‘parameterisations’ of the models where their effects are accounted-for in approximate form on a coarser model grid. Parameterisations are typically the result of intensive theoretical and observational studies and represent an additional level of detail within the climate model itself. Most climate projection models perform reasonably well, and it is generally accepted that there is no single ‘best’ climate model or subset of models. The relatively coarse scale of the grid used by the global models limit their ability to represent some important regional and local-scale features of climate though model resolution has improved over time. A technique known as down-scaling can be used to allow fine resolution regional models to be embedded within a global model.

Climate change models from the Coupled Model Inter Comparison Project Phase 5 (CMIP5) (49) were selected as candidate models for this study. These provide improvements on previous climate models which included an increase in horizontal and vertical resolution of predictions, an improved representation of processes within the climate system (i.e., aerosol-cloud interactions) and use of a larger number of ensemble (i.e., model contributor) members. For this study, we selected the ‘worst’ case RCP scenario in terms of temperature rise: CanESM2 model RCP 8.5. Daily average temperature data for the CanESM2 model RCP 8.5 scenario at a resolution of 0.05 × 0.05 decimal degrees (approximately 5 km × 5 km) were provided by the Australian Climate Futures (ACF) decision-support tool (49). ACF provides Australian climate projection data for 14 future time periods (from 2025 to 2090 in 5-yearly increments) and four scenarios of greenhouse gas concentrations (RCP 2.6, RCP 4.5, RCP 6.0 and RCP 8.5) using the latest CMIP5 models (49). Southern hemisphere summer (1 January) and winter (1 July) incursions of bluetongue were simulated for each of the two selected study areas (Figure 1).

Simulations were run in two modes. The first was with direct (i.e., farm-to-farm, farm-to-saleyard and saleyard-to-farm) animal movements disabled; the second was with direct animal movements enabled with movement frequencies and distances by herd type the same as that used for the AADIS model of foot and mouth disease (40). Our rationale for this approach was to enhance understanding of how animal movements between premises contribute to the predicted size of an outbreak, focusing on indirect spread mechanisms rather than specifying the distinct pathways of transmission. In addition, our study aimed to simulate the spread of disease without control measures to understand the potential extent of an outbreak under unmanaged conditions. This approach helps in assessing the maximum possible outbreak size, acknowledging that in a real-world scenario, especially for diseases like bluetongue, effective control strategies would be implemented, reducing animal movements and outbreak size. Thus, our findings represent a hypothetical scenario of unchecked spread, underscoring the critical role of timely detection and control in disease management. In summary, a total of 24 outbreak simulations were run: mid-summer and mid-winter incursions for each study area for each year (2015, 2025 and 2035) with direct movements first disabled, and then enabled for each. AADIS was run for 365 simulation days with 100 iterations used for direct movement-disabled simulations and for the direct movement-enabled simulations. The results of each of the 24 simulated outbreaks are described in terms of the total number of herds predicted to become infected, the size of the outbreak area (as defined by contour lines delineating 95% of infected herd densities, across all iterations) and the within-herd incidence risk for herds in which outbreaks of bluetongue occurred. Counts of the predicted number of infected herds across scenarios were compared using a two-tailed Welch (unequal variance) t test.

## Results

Minimum, mean and maximum daily temperatures for North Queensland and Northern New South Wales study areas are presented in Table 1 for 2015, 2025 and 2035, based on the CanESM2 model RCP 8.5 emission scenario. Note that our projections for 2025, developed in 2019, are now nearing the test of time as we approach the target year. The proximity of this date allows us to closely monitor unfolding events and trends, providing an opportunity to assess the accuracy of our predictions in real-time. These data are presented as line plots in Figure 2. Climate projections show an increase in mean and maximum average daily temperature for 2025 and 2035, compared with 2015. Maximum daily temperatures were predicted to be 1.9 ^○^C and 1.1 ^○^C higher in 2025 and 2035 (respectively) in the North Queensland study area, compared with 2015. In Northern New South Wales, minimum daily temperatures for 2025 and 2035 were predicted to be 0.8 ^○^C and 1.3 ^○^C higher (respectively), compared with 2015. The predicted densities of adult *Culicoides* spp. are presented as line plots in Figure 3 for the North Queensland and Northern New South Wales study areas for 2015, 2025 and 2035. While the predictions for 2015, 2025, and 2035 indicate a consistent trend in the abundance and activity of Culicoides for each study area, nuanced differences between the years were observed. Scenarios in North Queensland for 2025 and 2035 results showed relatively lower vector populations in winter compared with 2015. In Northern New South Wales, midge population predictions for 2025 and 2035 were higher and re-established quicker after winter than the predictions for 2015. Descriptive statistics of the number of herds predicted to be bluetongue positive by AADIS after 365 days following incursions of BTV into single herds in North Queensland and Northern New South Wales in mid-summer (1 January) and mid-winter (1 July) with direct animal movements disabled for 2015, 2025 and 2035 are shown in Table 2. Numbers of infected herds for the animal movement enabled scenarios are shown in Table 3. Maps of the North Queensland study area showing the point location of herds predicted by AADIS to become bluetongue positive following an incursion of BTV into a single herd in mid-summer (1 January) in 2015, 2025 and 2035 with direct animal movements disabled are shown in Figure 4. The same plots for the mid-winter North Queensland incursions are shown in Figure 5. Figures 6 and 7 show the same data for the Northern New South Wales study area. Box and whisker plots showing the distribution of the number of herds predicted to become bluetongue positive for the North Queensland and Northern New South Wales study areas, by season and year with direct animal movements disabled are shown in Figure 8.

**Figure 2:**
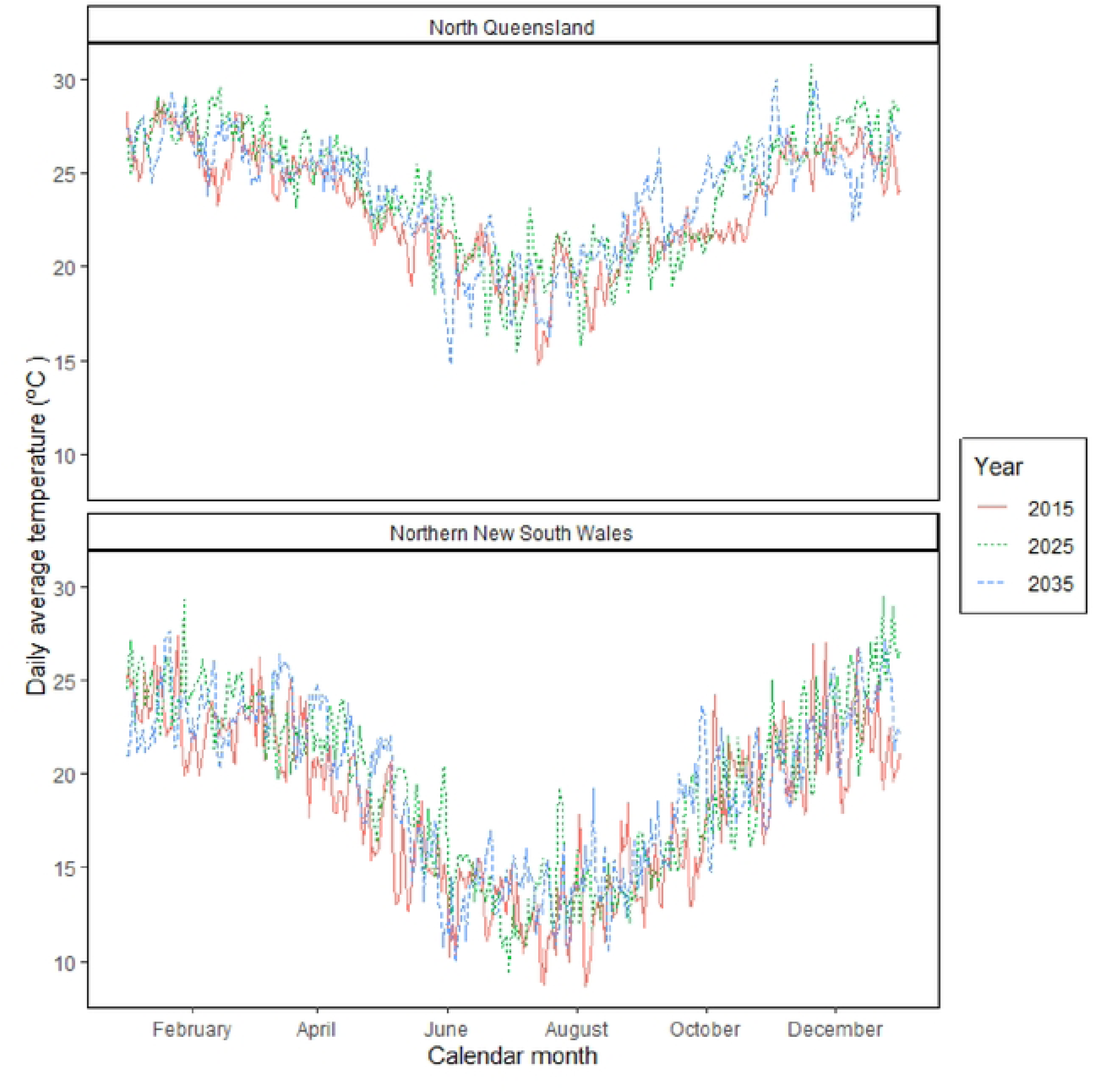
The effect of climate change on the spread of bluetongue in Australian livestock. Line plots showing average daily temperature as a function of day of the year for 2015, 2025 and 2035 for the North Queensland and Northern New South Wales study areas. The 2025 and 2035 scenarios are based on climate predictions from the CanESM2 model RCP 8.5 emission scenario.

**Figure 3:**
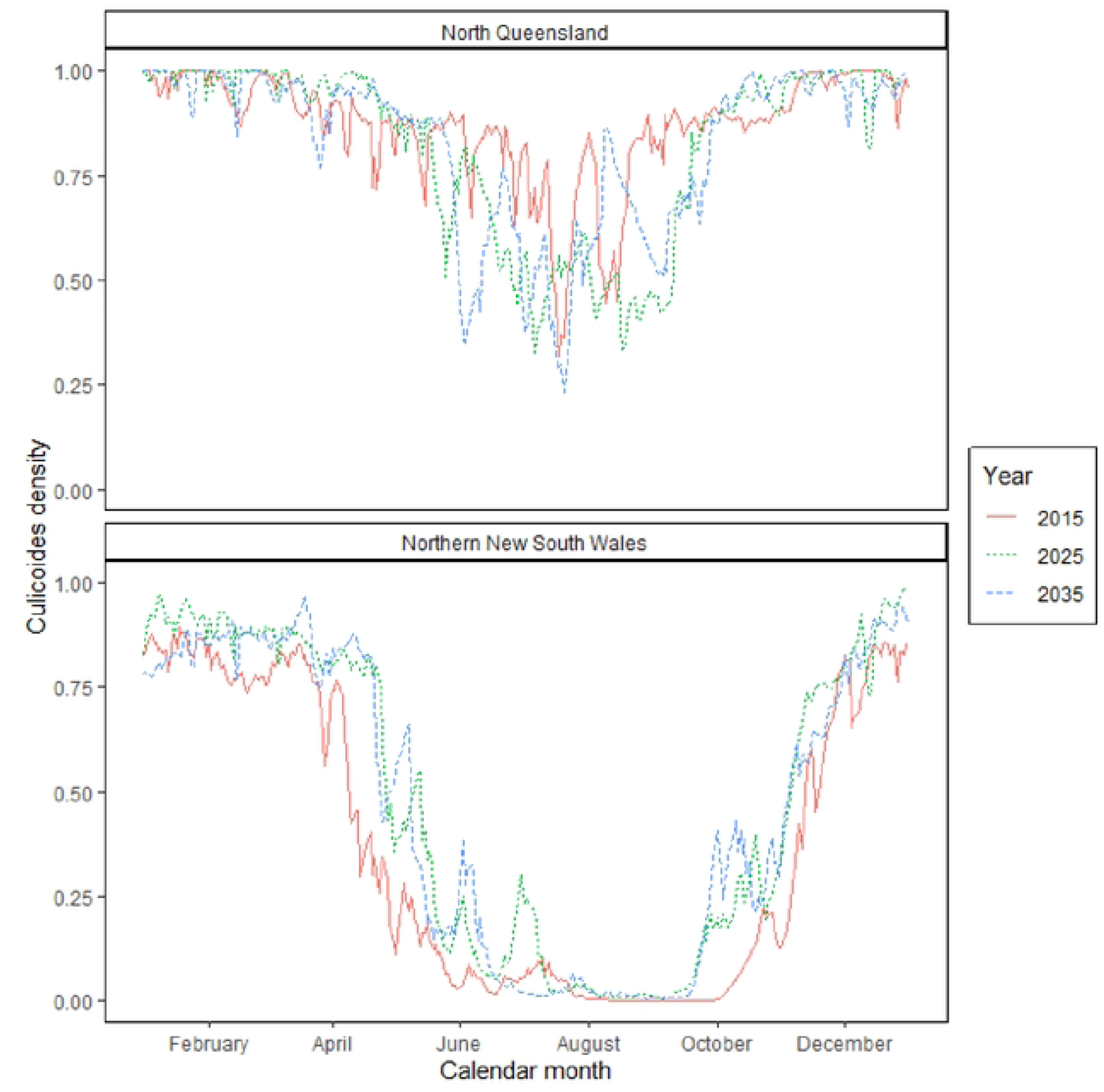
The effect of climate change on the spread of bluetongue in Australian livestock. Line plots showing average daily *Culicoides* spp. population density as a function of day of the year for 2015, 2025 and 2035 for the North Queensland and Northern New South Wales study areas. The 2025 and 2035 scenarios are based on climate predictions from the CanESM2 model RCP 8.5 emission scenario.

**Figure 4:**
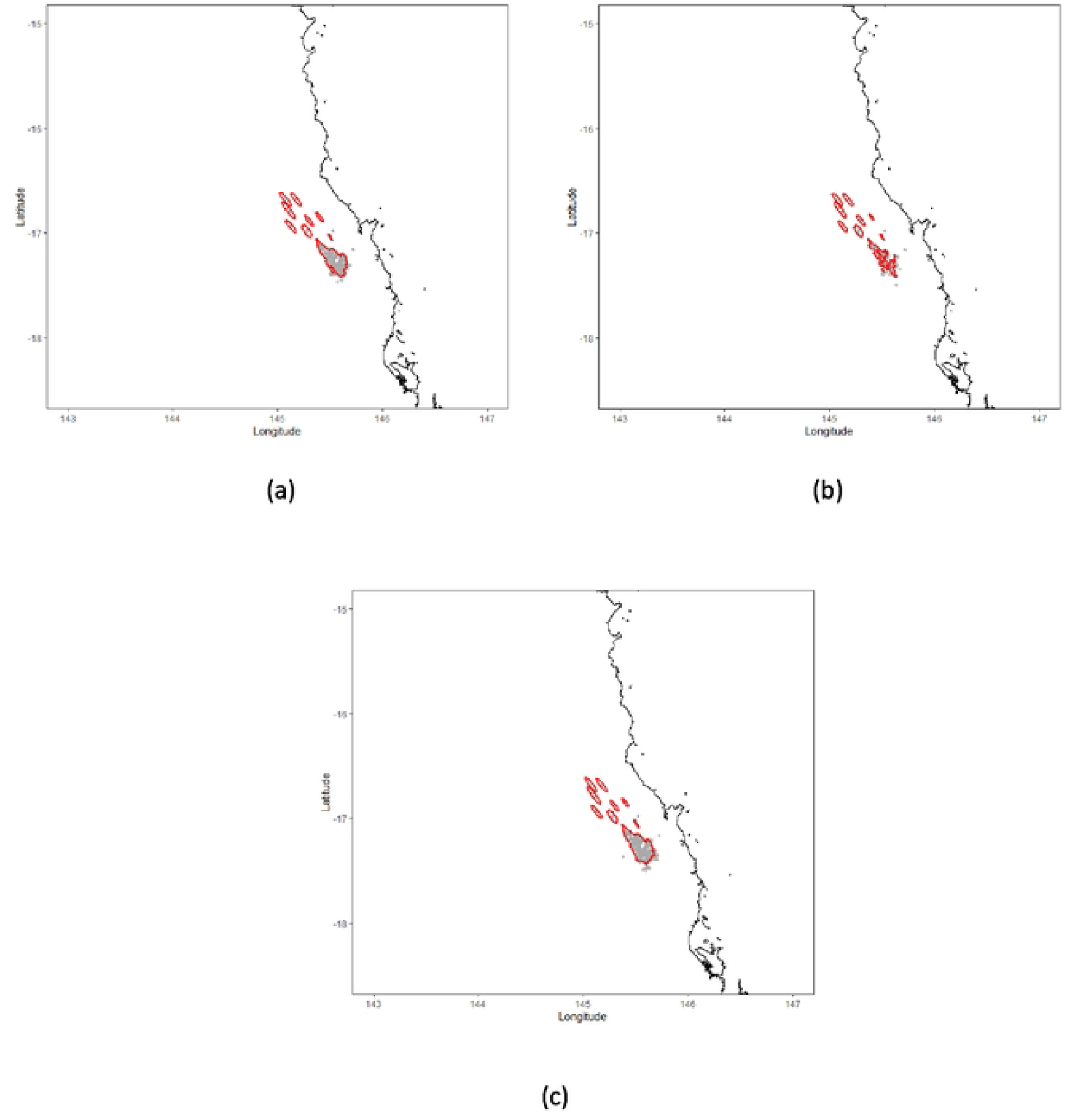
The effect of climate change on the spread of bluetongue in Australian livestock. Map of North Queensland study area showing the point location of herds predicted to become bluetongue positive by AADIS following incursion of BTV into a single herd in mid-summer (1 January) with direct animals movements disabled for: (a) 2015; (b) 2025; and (c) 2035. Contour lines delineate areas where the predicted number of bluetongue positive herds was greater than 95 per square kilometre. The 2025 and 2035 simulations are based on climate predictions from the CanESM2 model RCP 8.5 emission scenario. Predictions are based on 100 simulations of the AADIS bluetongue model, as described in the text.

**Figure 5:**
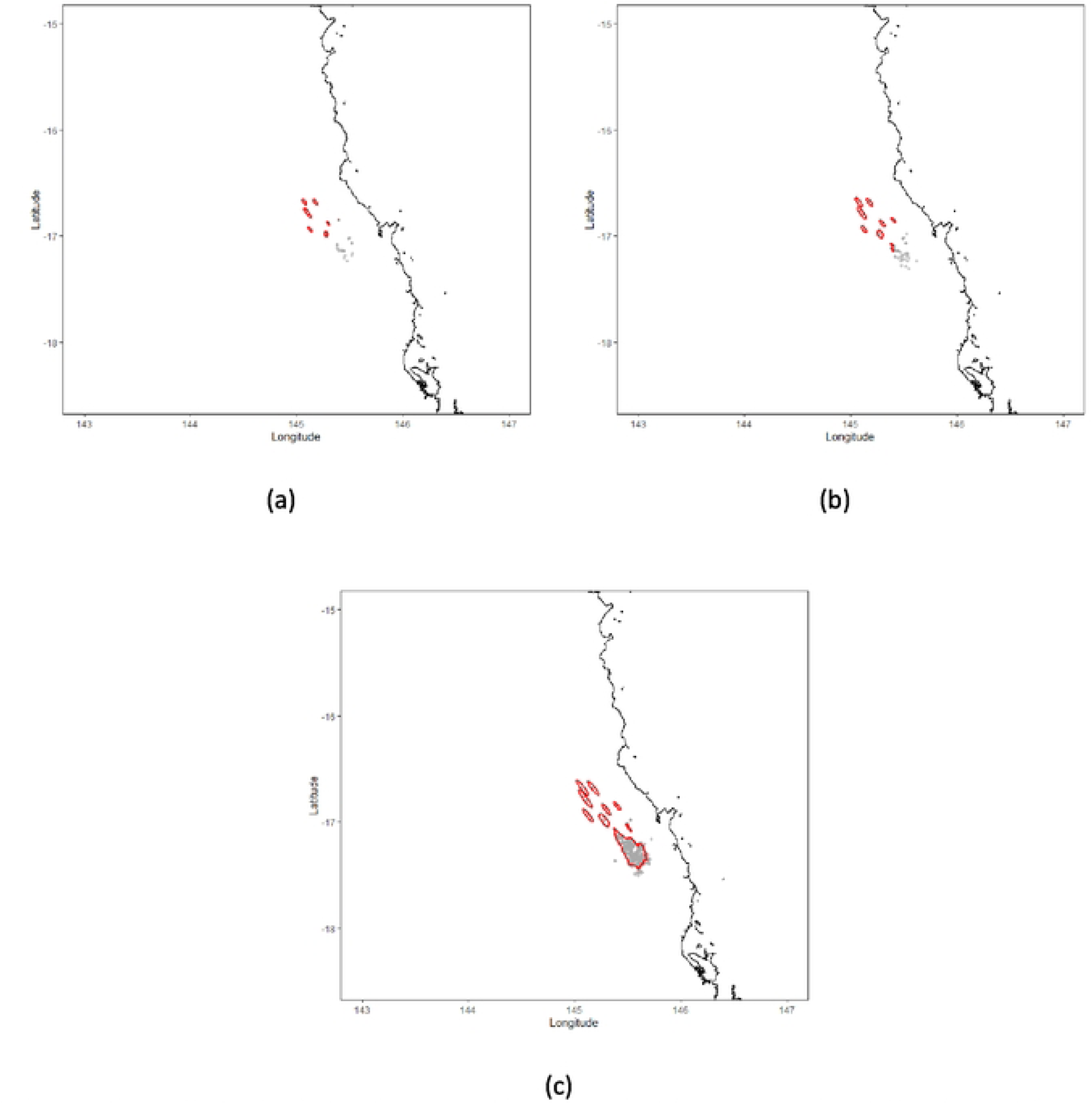
The effect of climate change on the spread of bluetongue in Australian livestock. Map of North Queensland study area showing the point location of herds predicted to become bluetongue positive by AADIS following incursion of BTV into a single herd in mid-winter (1 July) with direct animals movements disabled for: (a) 2015; (b) 2025; and (c) 2035. Contour lines delineate areas where the predicted number of bluetongue positive herds was greater than 95 per square kilometre. The 2025 and 2035 simulations are based on climate predictions from the CanESM2 model RCP 8.5 emission scenario. Predictions are based on 100 simulations of the AADIS bluetongue model, as described in the text.

**Figure 6:**
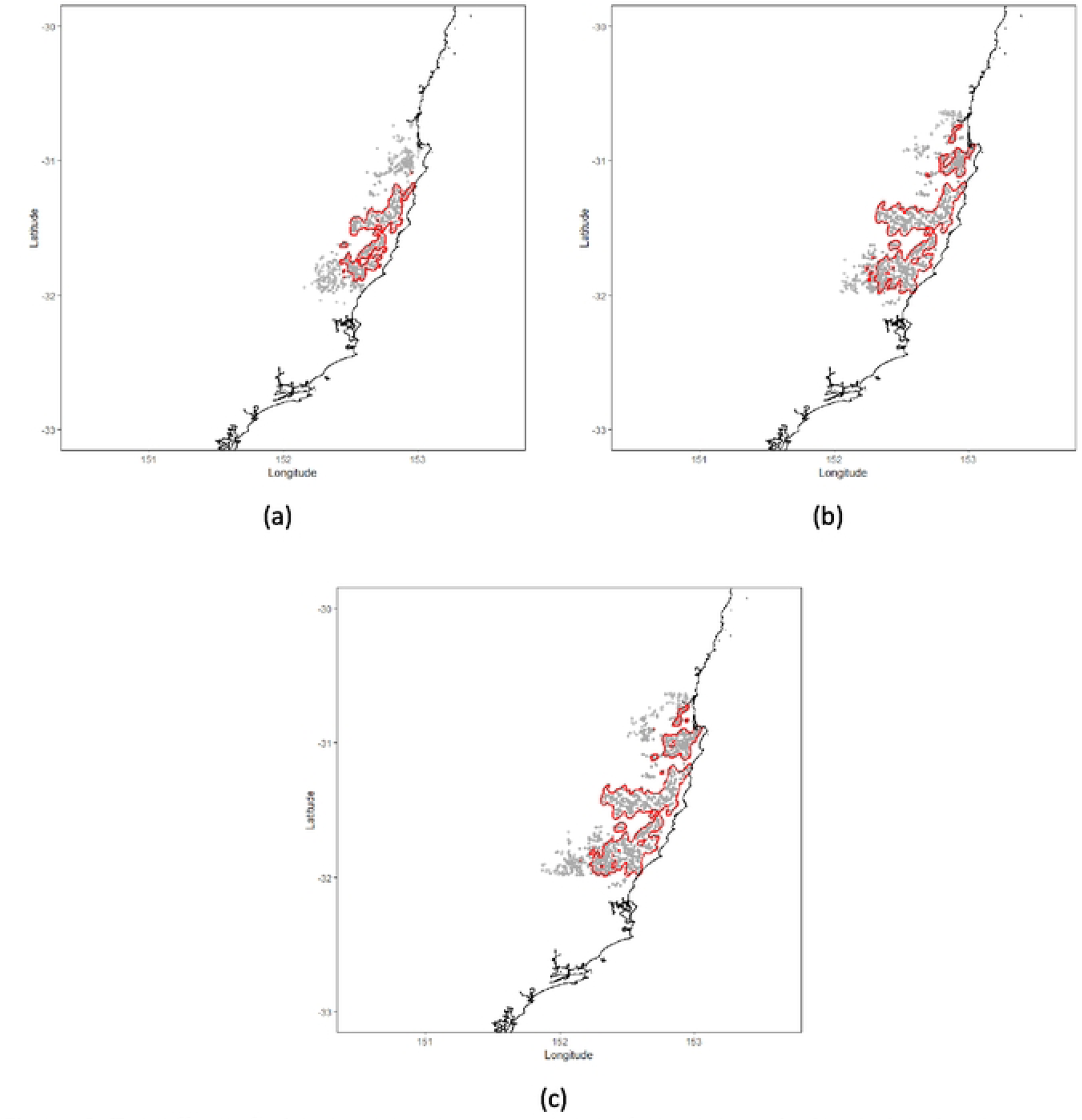
The effect of climate change on the spread of bluetongue in Australian livestock. Map of the Northern New South Wales study area showing the point location of herds predicted to become bluetongue positive by AADIS following incursion of BTV into a single herd in mid-summer (1 January) with direct animals movements disabled for: (a) 2015; (b) 2025; and (c) 2035. Contour lines delineate areas where the predicted number of bluetongue positive herds was greater than 95 per square kilometre. The 2025 and 2035 simulations are based on climate predictions from the CanESM2 model RCP 8.5 emission scenario. Predictions are based on 100 simulations of the AADIS bluetongue model, as described in the text.

**Figure 7:**
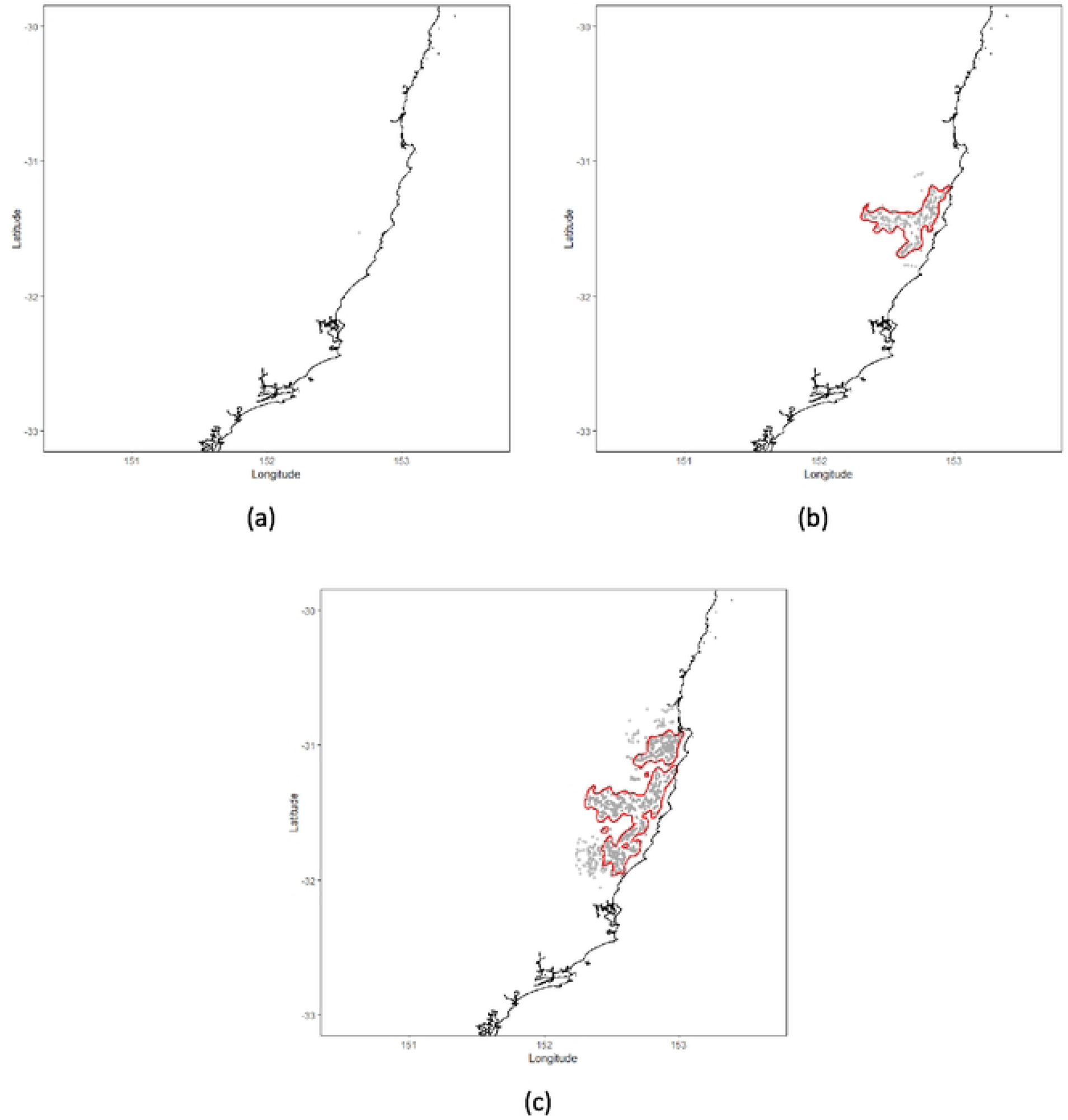
The effect of climate change on the spread of bluetongue in Australian livestock. Map of the Northern New South Wales study area showing the point location of herds predicted to become bluetongue positive by AADIS following incursion of BTV into a single herd in mid-winter (1 July) with direct animals movements disabled for: (a) 2015; (b) 2025; and (c) 2035. Contour lines delineate areas where the predicted number of bluetongue positive herds was greater than 95 per square kilometre. The 2025 and 2035 simulations are based on climate predictions from the CanESM2 model RCP 8.5 emission scenario. Predictions are based on 100 simulations of the AADIS bluetongue model, as described in the text.

**Figure 8:**
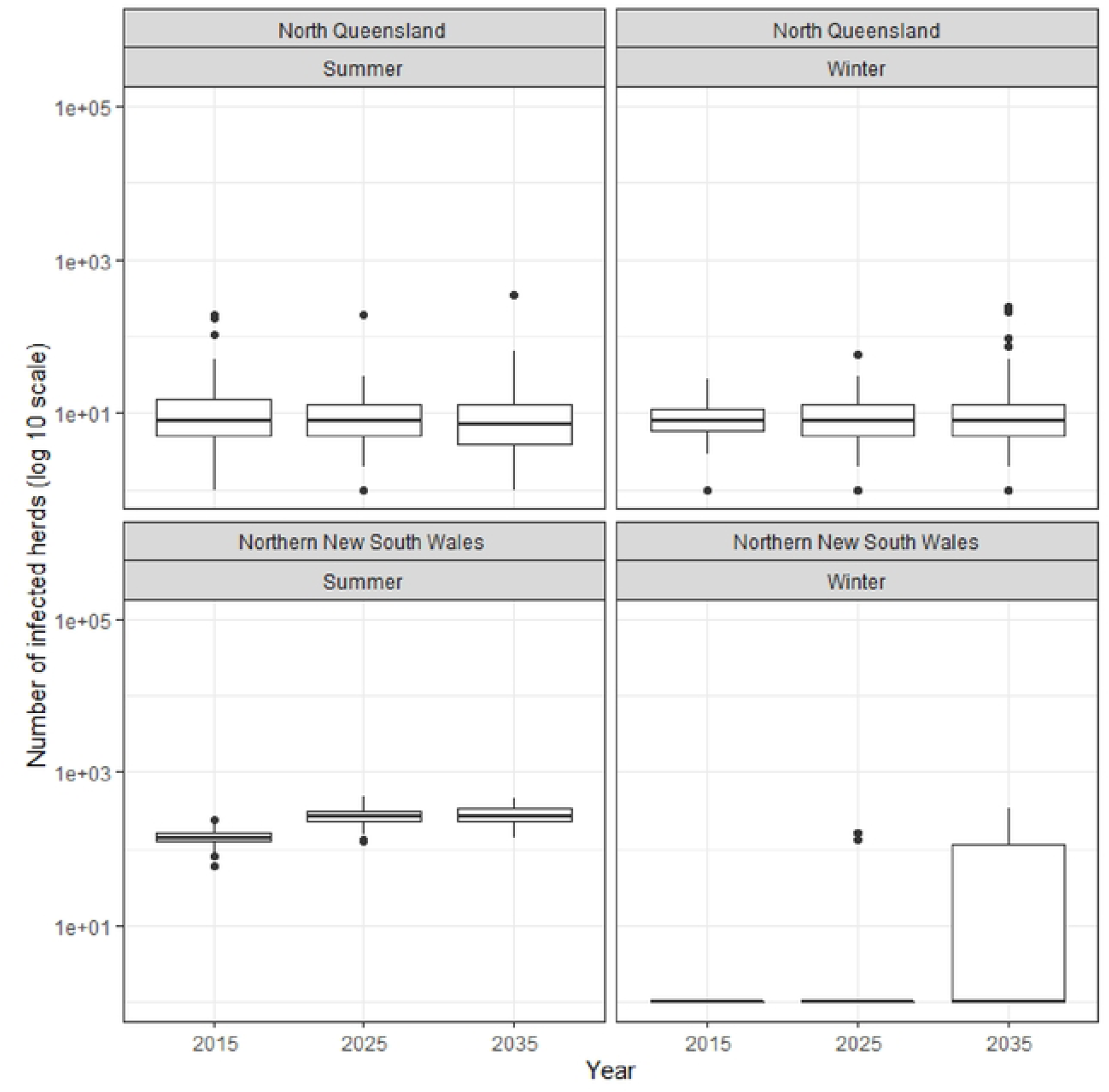
The effect of climate change on the spread of bluetongue in Australian livestock. Box and whisker plots showing the distribution of the number of herds predicted to become bluetongue positive by AADIS following incursion of BTV into single herds in North Queensland and Northern New South Wales in mid-summer (1 January) and mid-winter (1 July) with direct animals movements disabled for 2015, 2025 and 2035. The 2025 and 2035 simulations are based on climate predictions from the CanESM2 model RCP 8.5 emission scenario. Predictions are based on 100 simulations of the AADIS bluetongue model, as described in the text.

**Table 1:** Minimum, mean and maximum actual (2015) and predicted (2025 and 2035) average temperatures for the North Queensland and Northern New South Wales study areas described in the text. Numbers in brackets equal the change in temperature relative to 2015. Predicted temperatures are based on the CanESM2 model RCP 8.5 emission scenario.

| YEAR | AVERAGE DAILY TEMPERATURE (°C) |  |  |
| --- | --- | --- | --- |
|  | Minimum | Mean | Maximum |
| **NORTH QUEENSLAND:** |  |  |  |
| **2015** | 14.8 | 32.2 | 28.9 |
| **2025** | 15.3 (+0.5) | 24.1 (+0.9) | 30.8 (+1.9) |
| **2035** | 14.7 (-0.1) | 23.9 (+0.7) | 30.0 (+1.1) |
| **NORTHERN NEW SOUTH WALES:** |  |  |  |
| **2015** | 8.6 | 18.2 | 27.4 |
| **2025** | 9.5 (+0.8) | 19.4 (+1.2) | 29.5 (+2.1) |
| **2035** | 9.9 (+1.3) | 19.3 (+1.0) | 27.6 (+0.2) |

**Table 2:**
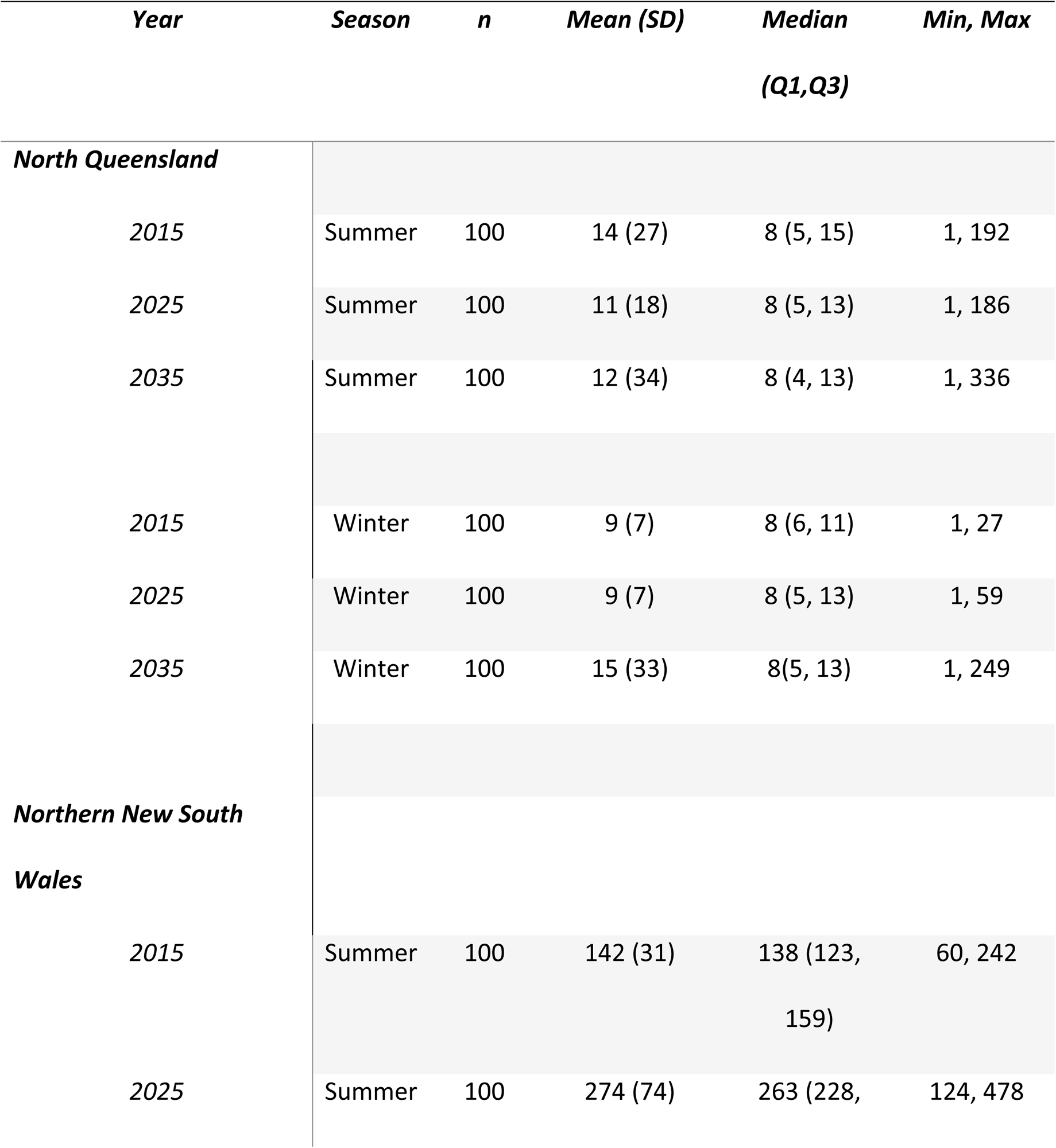
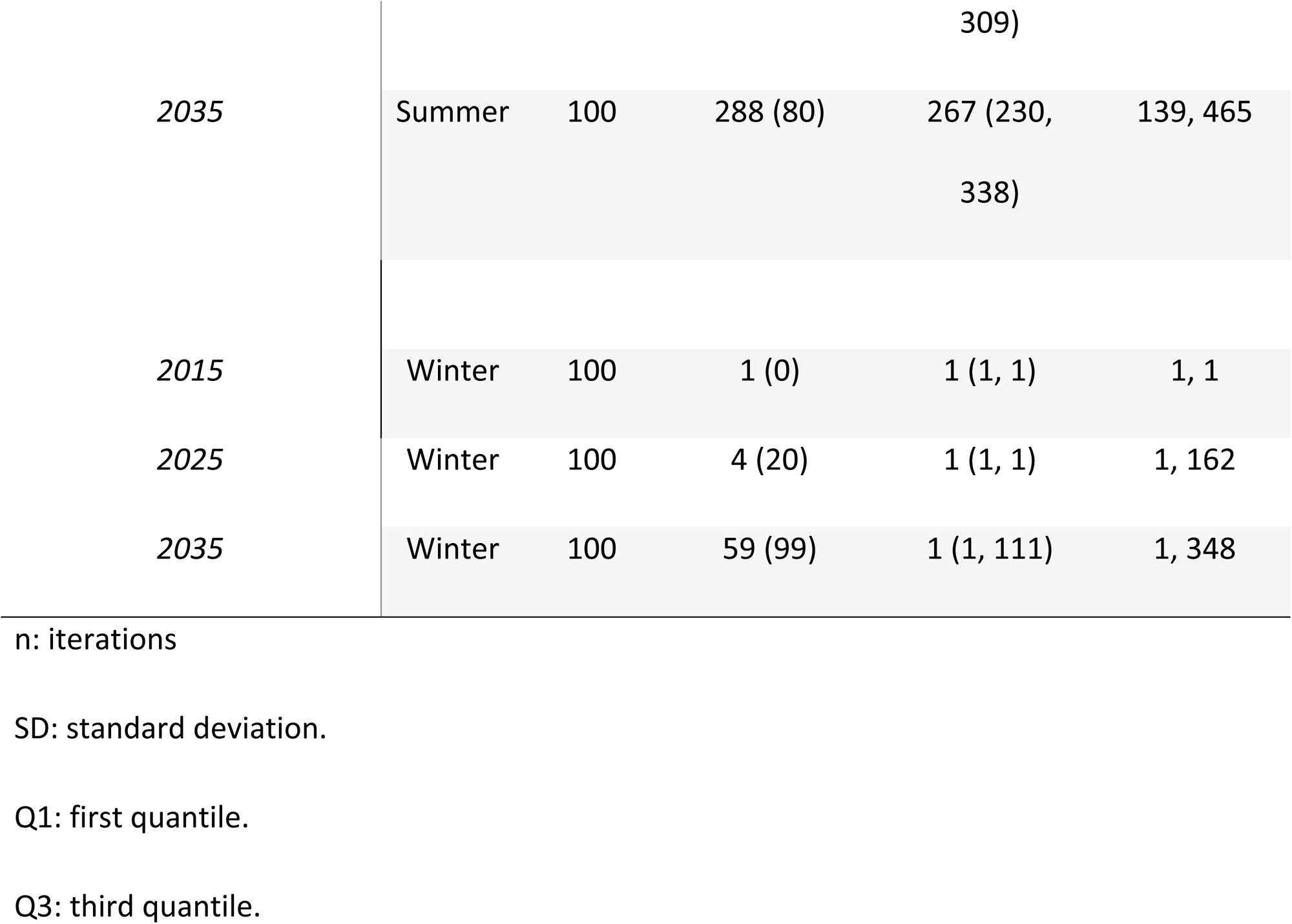
Descriptive statistics of the number of herds predicted by AADIS to be bluetongue positive after 365 days following incursions of BTV into single herds in North Queensland and Northern New South Wales in midsummer (1 January) and mid-winter (1 July) with direct animal movements disabled for 2015, 2025 and 2035. The 2025 and 2035 simulations are based on climate predictions from the CanESM2 model RCP 8.5 emission scenario.

**Table 3:**
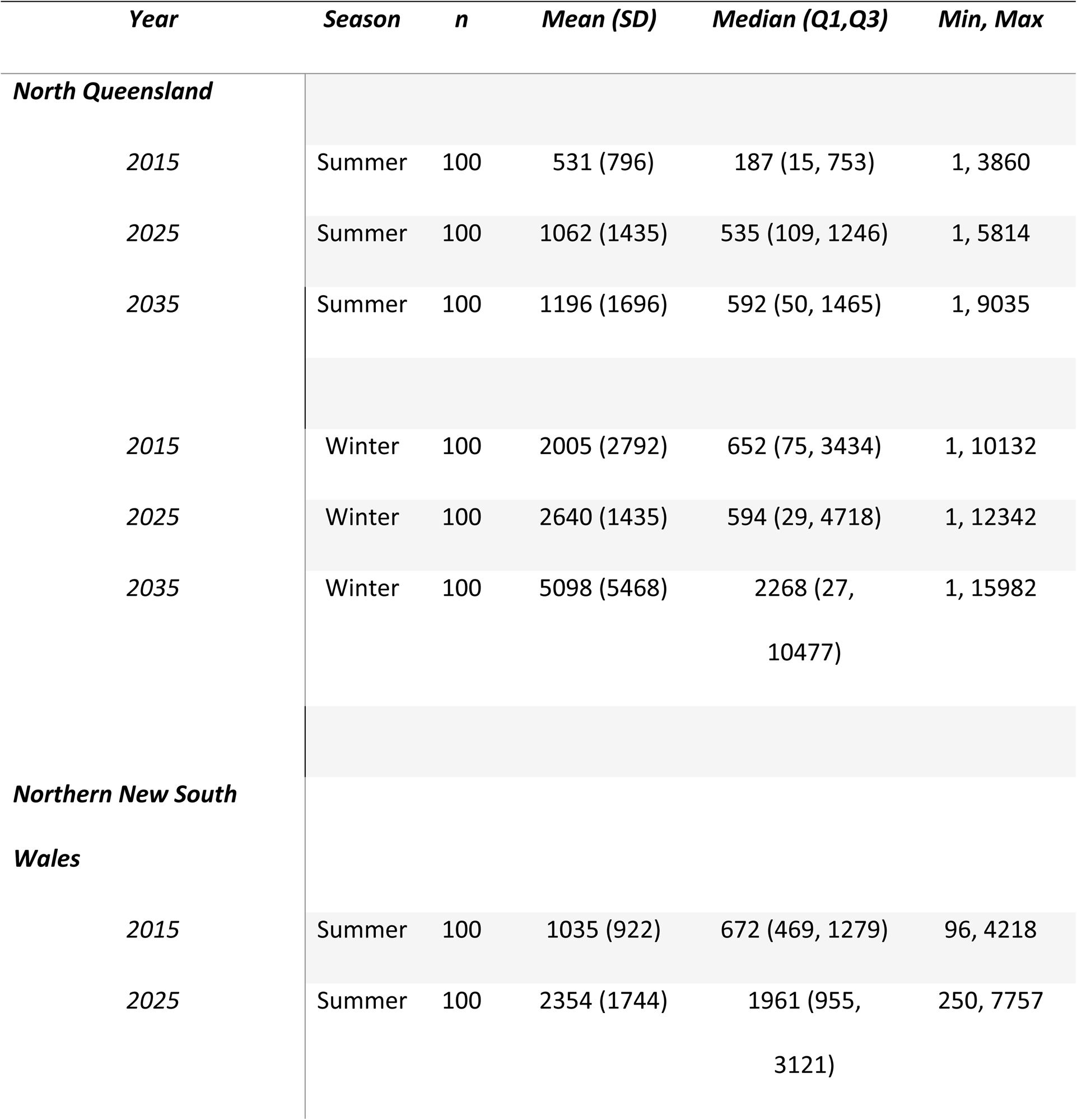
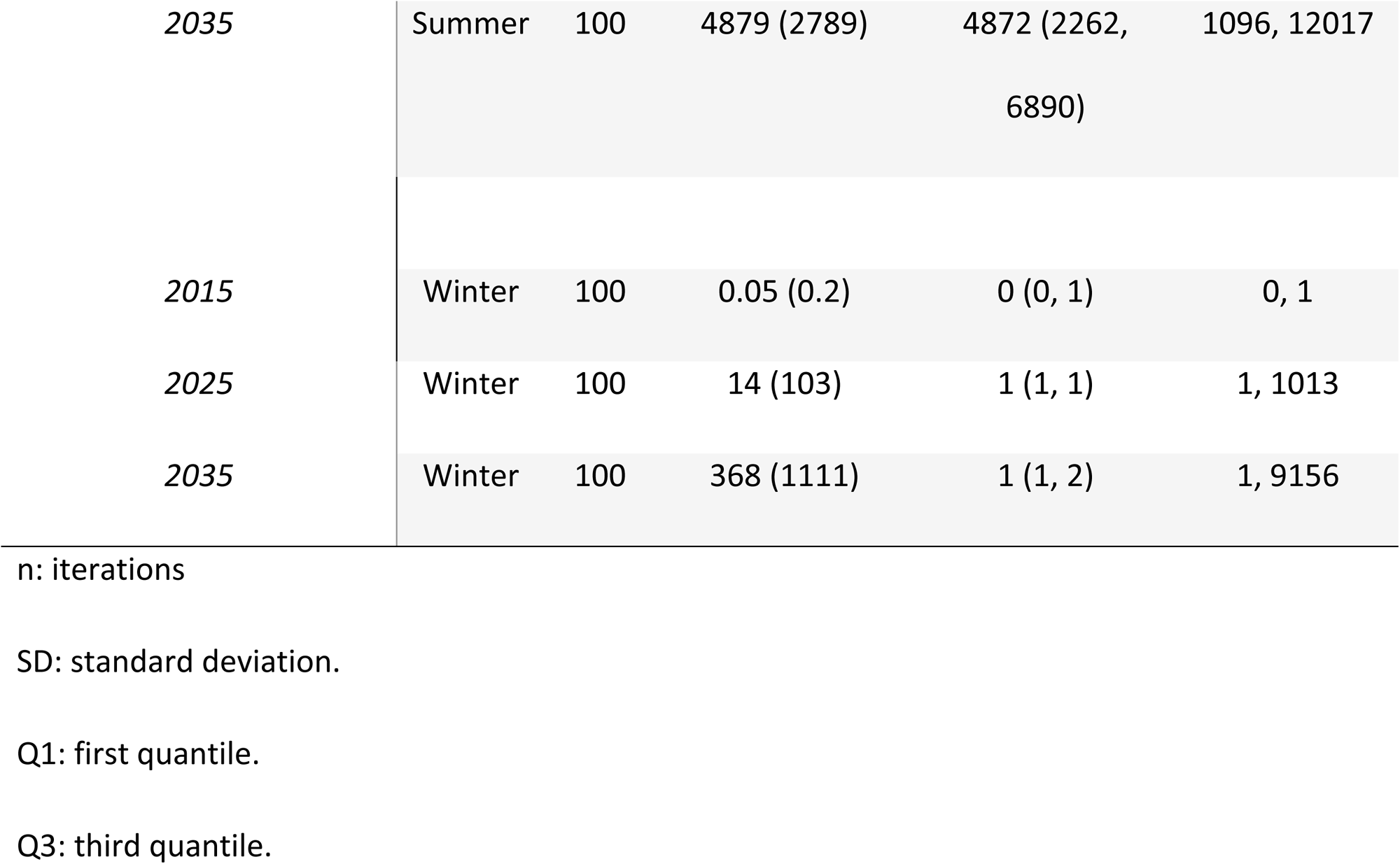
Descriptive statistics of the number of herds predicted by AADIS to be bluetongue positive after 365 days following incursions of BTV into single herds in North Queensland and Northern New South Wales in midsummer (1 January) and mid-winter (1 July) with direct animal movements enabled for 2015, 2025 and 2035. The 2025 and 2035 simulations are based on climate predictions from the CanESM2 model RCP 8.5 emission scenario.

Maps of Australia showing the point location of herds predicted to become bluetongue positive by AADIS following incursion of BTV into a single herd in North Queensland in mid-summer (1 January) in 2015, 2025 and 2035 with direct animal movements enabled are shown in Figure 9. The same plots for the mid-winter North Queensland incursions are shown in Figure 10. Similar to the direct movement disabled scenarios, in Figures 9 to 12 contour lines delineate areas where the predicted number of bluetongue positive herds was greater than 95 per square km. Figures 11 and 12 show the same data for the Northern New South Wales study area. Figure 13 are box and whisker plots showing the distribution of the number of herds predicted to become bluetongue positive for the North Queensland and Northern New South Wales study areas, by season and year with direct animal movements enabled.

**Figure 9:**
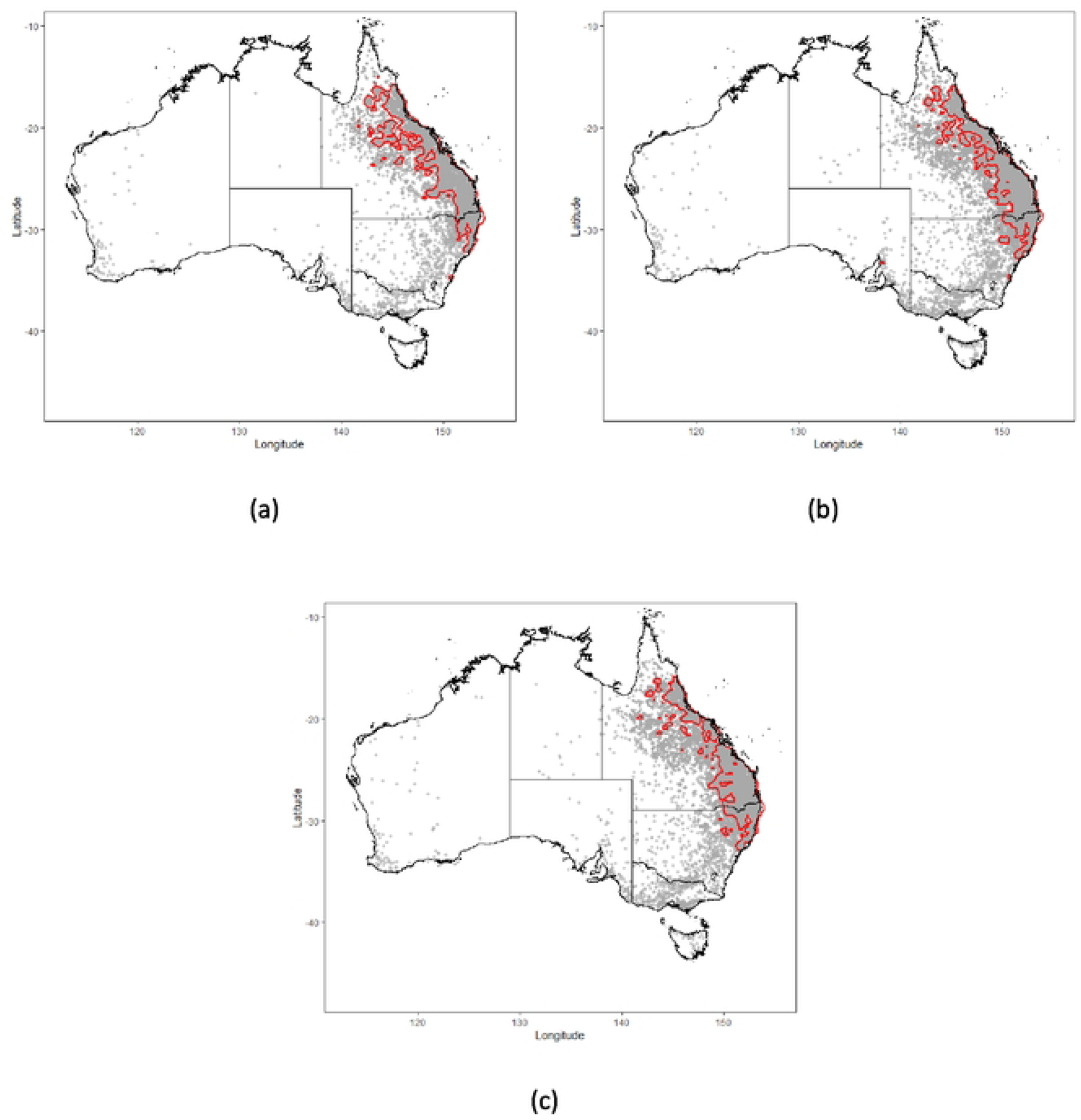
The effect of climate change on the spread of bluetongue in Australian livestock. Map of Australia showing the point location of herds predicted to become bluetongue positive by AADIS following incursion of BTV into a single herd in North Queensland in mid-summer (1 January) with direct animals movements enabled for: (a) 2015; (b) 2025; and (c) 2035. Contour lines delineate areas where the predicted number of bluetongue positive herds was greater than 95 per square kilometre. The 2025 and 2035 simulations are based on climate predictions from the CanESM2 model RCP 8.5 emission scenario. Predictions are based on 50 simulations of the AADIS bluetongue model, as described in the text.

**Figure 10:**
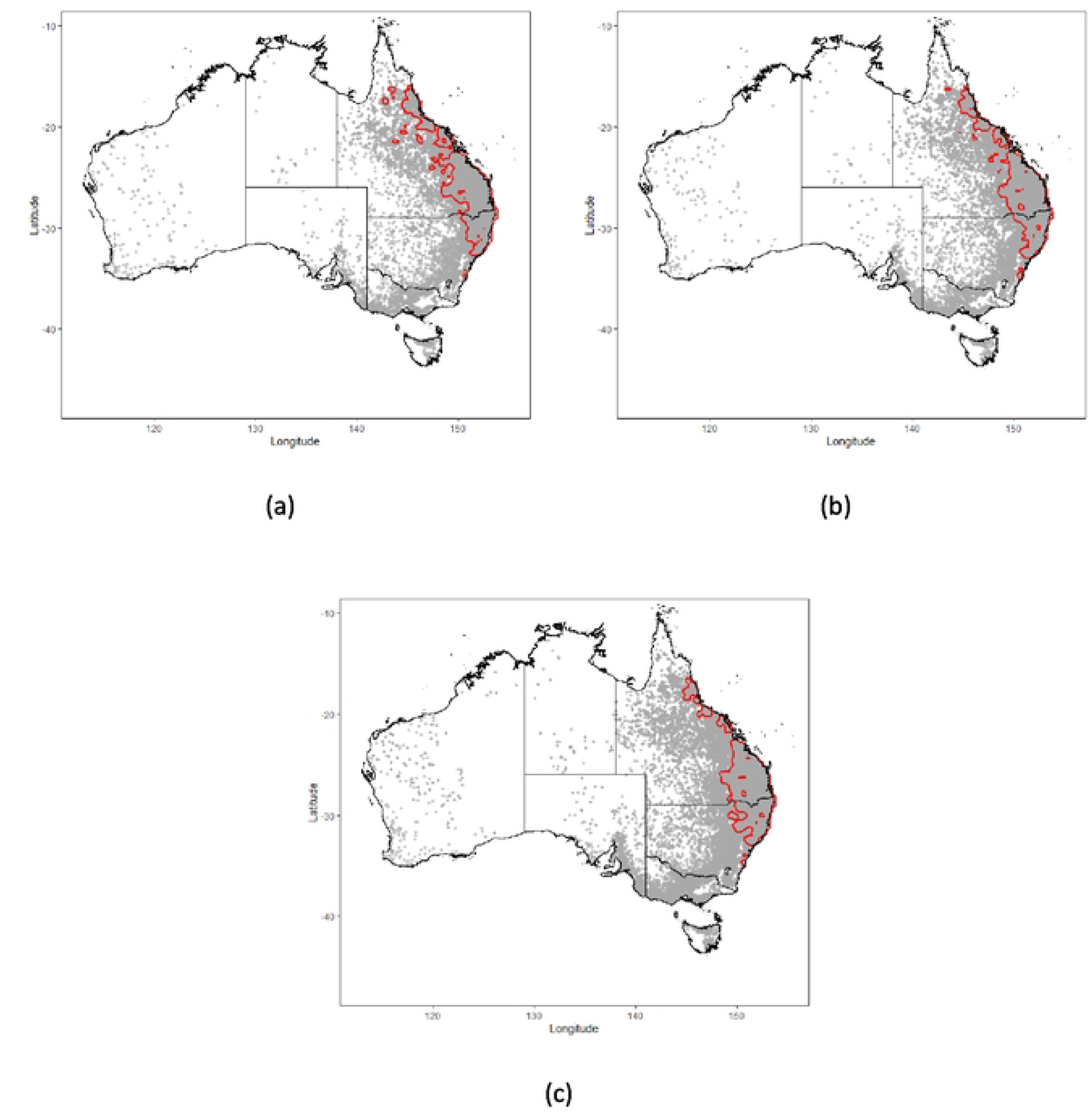
The effect of climate change on the spread of bluetongue in Australian livestock. Map of Australia showing the point location of herds predicted to become bluetongue positive by AADIS following incursion of BTV into a single herd in North Queensland in mid-winter (1 July) with direct animals movements enabled for: (a) 2015; (b) 2025; and (c) 2035. Contour lines delineate areas where the predicted number of bluetongue positive herds was greater than 95 per square kilometre. The 2025 and 2035 simulations are based on climate predictions from the CanESM2 model RCP 8.5 emission scenario. Predictions are based on 50 simulations of the AADIS bluetongue model, as described in the text.

**Figure 11:**
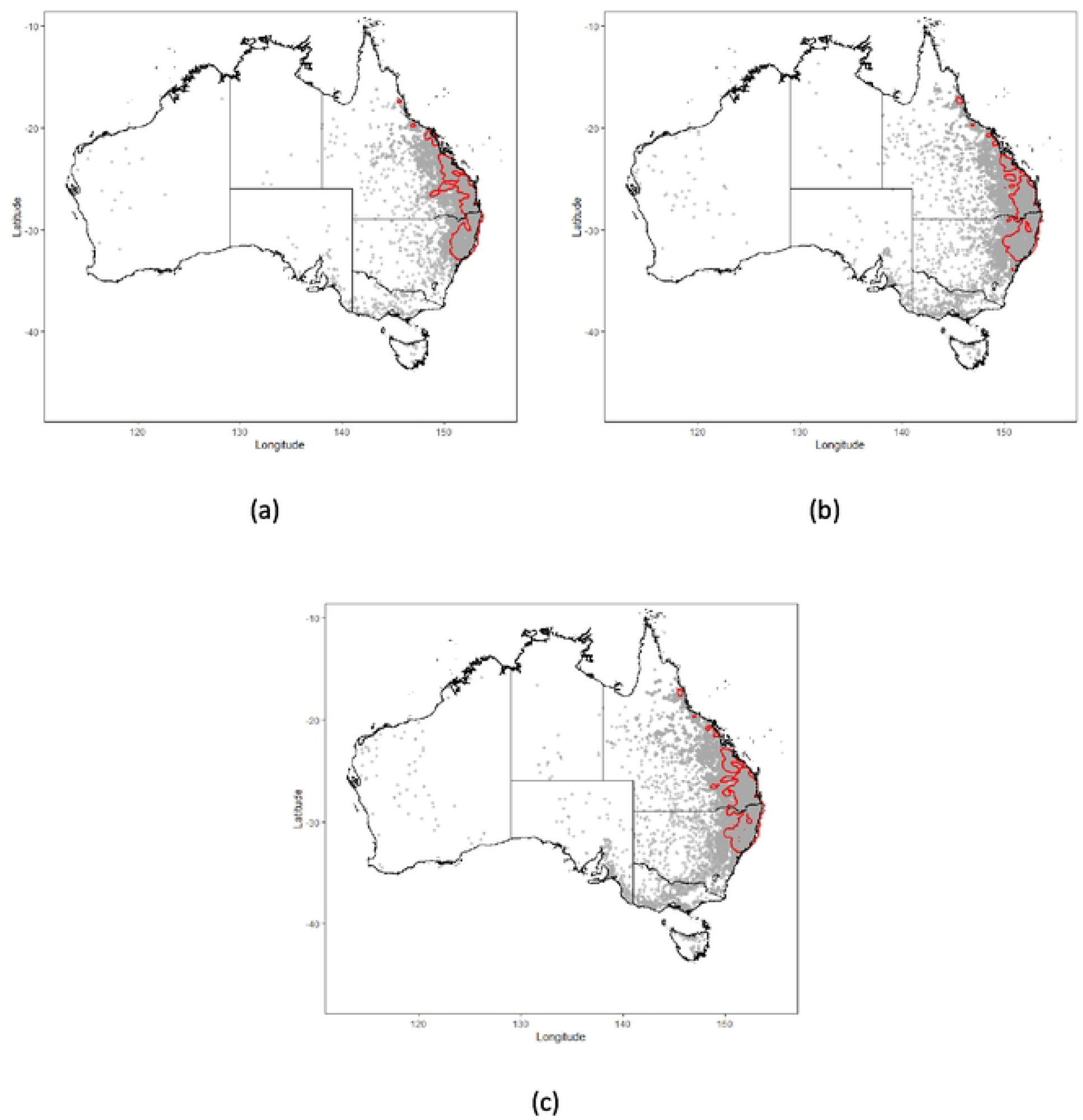
The effect of climate change on the spread of bluetongue in Australian livestock. Map of Australia showing the point location of herds predicted to become bluetongue positive by AADIS following incursion of BTV into a single herd in the Northern New South Wales in mid-summer (1 January) with direct animals movements enabled for: (a) 2015; (b) 2025; and (c) 2035. Contour lines delineate areas where the predicted number of bluetongue positive herds was greater than 95 per square kilometre. The 2025 and 2035 simulations are based on climate predictions from the CanESM2 model RCP 8.5 emission scenario. Predictions are based on 50 simulations of the AADIS bluetongue model, as described in the text.

**Figure 12:**
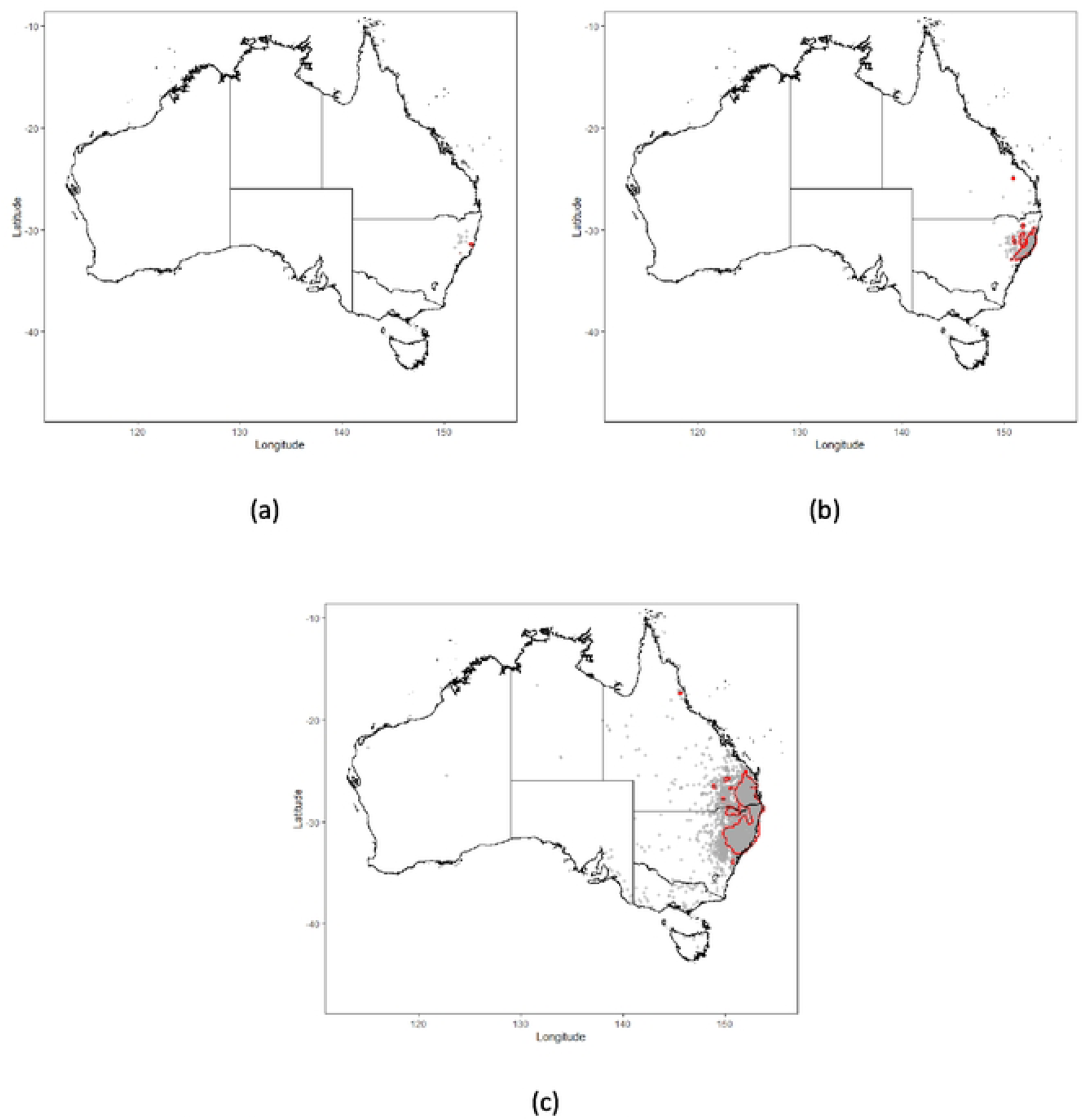
The effect of climate change on the spread of bluetongue in Australian livestock. Map of Australia showing the point location of herds predicted to become bluetongue positive by AADIS following incursion of BTV into a single herd in the Northern New South Wales in mid-winter (1 July) with direct animals movements enabled for: (a) 2015; (b) 2025; and (c) 2035. Contour lines delineate areas where the predicted number of bluetongue positive herds was greater than 95 per square kilometre. The 2025 and 2035 simulations are based on climate predictions from the CanESM2 model RCP 8.5 emission scenario. Predictions are based on 50 simulations of the AADIS bluetongue model, as described in the text.

**Figure 13:**
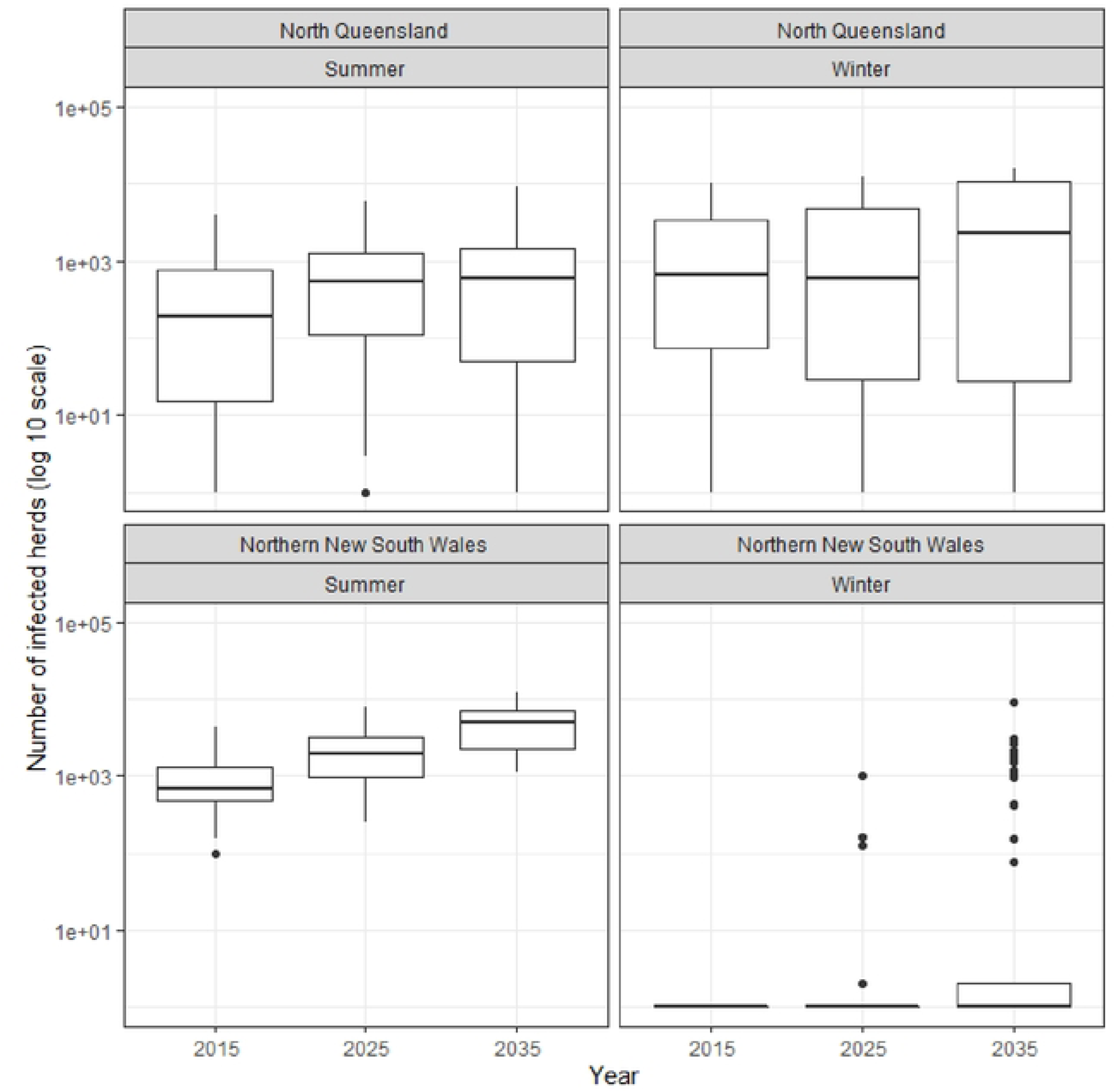
The effect of climate change on the spread of bluetongue in Australian livestock. Box and whisker plots showing the distribution of the number of herds predicted to become bluetongue positive by AADIS following incursion of BTV into single herds in North Queensland and Northern New South Wales in mid-summer (1 January) and mid-winter (1 July) with direct animals movements enabled for 2015, 2025 and 2035. The 2025 and 2035 simulations are based on climate predictions from the CanESM2 model RCP 8.5 emission scenario. Predictions are based on 50 simulations of the AADIS bluetongue model, as described in the text.

For the North Queensland, direct animal movement disabled scenarios our predictions showed there was little change in the median predicted number of bluetongue positive herds for the mid-summer and mid-winter incursions (Table 2, Figure 4 and Figure 5) although the variability of predicted outbreak size for the 2035 incursions (summer: minimum 1, maximum 336; winter: minimum 1, maximum 249) was greater than the variability of predicted outbreak size for the 2015 incursions (summer: minimum 1, maximum 192; winter: minimum 1, maximum 27). For Northern New South Wales there were moderate increases in predicted outbreak size for the summer incursions with a median of 142, 274 and 288 bluetongue positive herds for 2015, 2025 and 2035, respectively. Similar to North Queensland, the variability of predicted outbreak size for the 2035 incursions (summer: minimum 139, maximum 465; winter: minimum 1, maximum 348) was greater than the variability of predicted outbreak size for the 2015 incursions (summer: minimum 60, maximum 242; winter: minimum 1, maximum 1).

For the predicted number of bluetongue positive herds for the direct movement enabled scenarios were (predictably) much higher than the predicted number of bluetongue positive herds for the direct movement disabled scenarios (Table 3). Compared with the direct animal movement disabled scenarios, there was a clear trend in the increase in the predicted number of bluetongue positive herds as a function of simulation year for North Queensland (Figure 8). For Northern New South Wales this trend was not as marked, but as for the direct movement disabled scenarios, the predicted number of infected herds and the variability of predicted outbreak sizes for the 2035 incursions (summer: minimum 1096, maximum 12,017; winter: minimum 1, maximum 9156) was greater than the variability of predicted outbreak size for the 2015 incursions (summer: minimum 96, maximum 4218; winter: minimum 0, maximum 1). The predicted number of infected herds for the 2015 summer and 2035 summer incursions were significantly different (Welch t test statistic: - 9.49; *df* 54.42; P < 0.001). The predicted number of infected herds for the 2015 winter and 2035 winter incursions were significantly different (Welch t test statistic: -3.31; *df* 99; P < 0.001).

## Discussion

The aim of this study was to investigate the effect of climate change on potential BTV future spread in Australia. Estimates of the geographic distribution of *Culicoides* spp. across Australia (43) were combined with a model of bluetongue transmission (43) which was then implemented within an existing model of infectious animal disease (AADIS). Assuming the worst-case scenario for greenhouse gas emission (RCP 8.5) the key insight from our analysis is the modest increase in the predicted number of infected herds for summer BTV incursions into North Queensland and Northern New South Wales in 2025 and 2035, compared to 2015. This stability, observed even in scenarios without direct animal movements, reveals a relatively consistent pattern of BTV spread under short-term climate change projections. These findings, while focusing on the near future, emphasize that the changes occurred within a relatively short timeframe. It suggests that, over a longer period, such as 100 years, we could expect more pronounced changes, highlighting the potential for significant shifts in disease dynamics under extended climate change scenarios (Table 2). In our study, winter incursions in Northern New South Wales (NSW) for 2025 and 2035 showed a substantial increase in bluetongue-positive herds, compared with those in 2015. This region, distinct from Queensland (Qld) due to its climatic and geographical characteristics, presents unique conditions that influence disease spread. Northern NSW, with its cooler climate and varied topography, contrasts with Queensland’s warmer and more consistent climate. These differences can affect vector activity and disease transmission dynamics, making NSW a critical area for observing bluetongue trends over time. Our findings underscore the importance of understanding regional nuances in disease spread, particularly for international readers unfamiliar with Australian geography and climate variations. The recovery of Culicoides spp. post-winter involves a resurgence in population as environmental conditions improve. Warmer temperatures and increased moisture after winter end the dormancy period, triggering a rapid increase in activity and reproduction. Climate change exacerbates this effect by causing earlier springs and milder winters, leading to faster and more pronounced recovery periods. This enhanced recovery contributes to the observed higher abundance of Culicoides spp., as detailed in Figure 3, where the link between climatic factors and population dynamics is explored. AADIS allowed us to develop a better understanding of the role of direct animal movements as a determinant of outbreak size (compare the predicted number of infected herds in Table 2 with the predicted number of infected herds in Table 3). Importantly, the variability in the predicted size of bluetongue outbreaks was greater for the 2025 and 2035 incursions, compared with 2015, highlighting the impact of future temperature projections on disease dynamics. This increased variability underscores the challenge of predicting outbreak magnitudes under changing climatic conditions, emphasizing the need for adaptable management strategies in the face of climate-induced uncertainty. When direct animal movements were enabled, there was a substantial increase in the number of infected herds for the 2025 and 2035 incursions, compared with 2015, with greater numbers of infected herds when incursions occurred during the winter (see Table 3 and Table 2). The most likely reason for this is that in North Queensland off-farm movement events are more frequent during the winter and early spring (May to November) period (Iglesias & East 2015). Seeding of viraemic cattle into areas with a competent insect vector population (particularly in animal dense areas) meant that a new host-vector-host cycle of transmission could be established, further increasing infected herd numbers (Figures 9 to 12).

Previous European studies using climate-based ecological niche or R_0_(T) models identified an increase in bluetongue risk over time (Samy & Peterson 2016, Brand & Keeling 2017) and a marked increase in herd-level bluetongue incidence risk in 2006, consistent with the onset of the European BTV serotype 8 outbreak (30). Although R_0_(T) models provide useful estimation of bluetongue transmission potential (with values of R_0_(T) >1 indicative of sustained virus transmission) such models are unable to predict which herds will become infected during an outbreak or which combination of disease control measures will be successful.

Climate change is anticipated to significantly affect the geographic distribution of Culicoides spp., the insect vector for bluetongue in Australia, by expanding the temperature range (13.5°C to 35°C) conducive to their survival (50). With climate change, a larger area of the country is expected to experience temperatures within this range for extended periods annually. Understanding the ecology of Culicoides brevitarsis reveals that favorable temperatures lead to a southward expansion of their distribution, while unfavorable conditions cause a retreat northward or a quiescence in their activity. In extreme cold, populations may die off, leading to a retraction of their distribution to warmer northern regions. Over time, climate change is likely to make southern regions more habitable for these vectors, pushing the endemicity line progressively southward and altering the landscape of disease risk in Australia. Furthermore, the tendency for *Culicoides* spp. midges to feed on mammalian hosts increases as a function of temperature, as does the length of time taken for *Culicoides* midges to become infectious once they have been exposed to BTV (13, 51). However, the anticipated increase in extreme temperature days during warmer seasons might decrease vector abundance by pushing conditions beyond the ideal range for Culicoides growth, thus reducing survival rates. Nevertheless, for NSW, projections for 2025 and 2035 indicate that the population may not experience as significant a decline during colder months. This resilience could be linked to a warming trend, leading to less severe cold periods and enabling the Culicoides population to sustain higher numbers throughout the year. These findings imply that in NSW, milder winter conditions due to climate change may mitigate the population’s seasonal reduction, maintaining a stable vector presence even in colder months. Environmental temperature is not the only factor that determines the changing risk of BTV outbreaks but the emergence of bluetongue in Europe supports the conjecture that climate change was a contributory factor (51). Modelling studies show that in recent decades Europe’s warming has allowed *C. imicola* to extend beyond its established geographic range (52). This conclusion was supported by our study where the size of the summer and winter outbreaks that occurred in Northern New South Wales were greater for the 2025 and 2035 incursions, compared with 2015. In a study of the effect of climate change on the global distribution of bluetongue Samy and Peterson (2016) predicted a marked increase in the geographic distribution of bluetongue for RCP 8.5 compared with RCP 2.6. Increases in vector abundance does not necessarily lead to increases in the size of predicted bluetongue outbreaks but instead depends on an interaction between herd and animal density, vector abundance and animal movement frequencies. Under the effect of climate change and in the event of an incursion of virulent bluetongue in Australia, early detection and rapid imposition of effective restrictions on animal movement provide the best opportunity for rapid disease control. The data presented in this study permits a somewhat more refined assessment of how climate change may alter bluetongue risk in Australia and allows bluetongue surveillance and bluetongue incursion response planning in the coming years to be adapted accordingly.

